# Informed and uninformed empirical therapy policies

**DOI:** 10.1101/629550

**Authors:** Nicolas Houy, Julien Flaig

## Abstract

We argue that a proper distinction must be made between informed and uninformed decision making when setting empirical therapy policies, as this allows to estimate the value of gathering more information and to set research priorities. We rely on the stochastic version of a compartmental model to describe the spread of an infecting organism in a health care facility, and the emergence and spread of resistance to two drugs. We focus on information and uncertainty regarding the parameters of this model. We consider a family of adaptive policies. In the uninformed setting, the best adaptive policy allows to reduce the average cumulative infected patient-days over two years by 39.3% (95% CI: 30.3% – 48.1%) compared to the combination therapy. Choosing empirical therapy policies while knowing the exact parameter values allows to further decrease the cumulative infected patient-days on average by 3.9% (95% CI: 2.1% – 5.8%). In our setting, the benefit of perfect information might be offset by increased drug consumption.

## 1 Introduction

Antimicrobials are known to select for drug resistance [17]. Their extensive and indispensable use, as well as patients’ vulnerability, make drug resistance a serious issue in health care facilities [9]. Since antimicrobial resistance is often associated with fitness costs, that is a lower ability of resistant organisms to survive and reproduce in the absence of treatment compared to their drug susceptible counterpart, strategies have been proposed to counteract the evolution of resistance by treating strains with different drugs in empirical therapies, either over time or simultaneously [18]. An empirical therapy policy specifies which antimicrobial is to be administered “by default” before the exact cause of an infection is known.^1^ Empirical therapy policies aimed at reducing the evolution and spread of antimicrobial resistance include alternating drugs (*drug cycling* or *metronomic therapies*, either following a fixed rotation schedule or adaptively, see [14]), and assigning drugs randomly to patients (*drug mixing*).

The respective merits of such policies have been debated and empirical evidence remains elusive. Theoretical studies have sought to compare the performance of empirical therapy policies by using compartmental models of infections and emergence of drug resistance in health care facilities at the between-host level. We refer to [6, 22] for reviews and perspectives on this topic. The emergence and spread of drug resistance is a nonlinear phenomenon that strongly depends on the parameters of the population dynamics in the health care facility and on the characteristics of the disease. These parameters, however, may be difficult or costly to estimate accurately, thus preventing the choice of an optimal empirical therapy policy in each given parameter setting.

The problem of making decisions under parameter uncertainty is ubiquitous in health care. Put in general terms, the decision maker needs to choose an alternative *a* in a set *A* of strategies so as to maximize a value function *V* (*a, ξ*), where *ξ* is a set of uncertain parameters. An alternative *a*_0_ is typically chosen that maximizes the expected value over possible realizations of *ξ*: *a*_0_ = argmax_*a∈A*_*E*_*ξ*_[*V* (*a, ξ*)].^2^ Notice that *V* (*a, ξ*) may itself be the expected outcome of a stochastic process that is not specified here. While maximizing the expected value by choosing *a*_0_, the decision maker forgoes alternatives that would perform better than *a*_0_ for some instances of *ξ*. Said differently, the performance of an alternative *a*_0_ chosen without perfect knowledge of *ξ* can go far from the optimum for each instance of *ξ*. The expected “loss” of the decision maker between choosing the best strategy on average over the possible realizations of *ξ* his best strategy for each realizations of *ξ* is then *E*_*ξ*_[max_*a∈A*_*V*(*a, ξ*) − *V*(*a*_0_, *ξ*)] = *E*_*ξ*_[max_*a∈A*_ *V*(*a, ξ*)] − max_*a∈A*_ *E*_*ξ*_[*V*(*a, ξ*)]. This quantity is called the expected value of perfect information (EVPI). The EVPI provides an upper bound on the cost one should be willing to pay to acquire more information on *ξ*. Value of information analyses have been proposed as an extension of sensitivity analyses [10, 11], and as a tool for research prioritization and clinical trial optimization [24]. Efficient estimation methods for variants of the EVPI have been developed until recently [12, 13].

While the distinction between uninformed best expected outcome and expected informed best outcome is a classic topic of the health economics literature [3], we believe that it deserves more attention than it has received to this day in investigations of optimal empirical therapy policies. Many authors in this literature used only point estimates of model parameters to investigate the effects of antimicrobial consumption on the emergence and spread of resistance [1, 2, 7, 8, 16]. Other authors performed sensitivity analyses of model parameters. In [20], for instance, the authors showed how the performance of each considered policy was affected by variations of each parameter when other parameters were set to their default value. This approach allows to compare the respective influences of parameters on the performance of a given policy, but it does not take account of the respective performances of policies in the parameter space. In [25], the authors considered six different scenarios and, for each of them, variations of each parameter over relevant values, all other parameters being set to their default value. Again, the chosen approach did not allow to explore the entire parameter space. Because of this, while the authors showed how the optimal policy changed with the parameters, they could not provide results in terms of the expected payoff of always choosing the best policy when the different parameter values are known. They gave results in terms of which policy outperformed the others “for most parameter settings”. This conclusion was clearly meant to provide insight to decision makers in the absence of information about parameter values. However it is not without ambiguity, and this for two reasons. First, policies that perform better in most cases do not necessarily maximize the expected payoff (policy rankings do not contain all relevant information about payoffs). Second, this way of presenting results may cause confusion as it seems to confound the two terms in the EVPI: the best policies are computed *for each parameter setting*, that is based on *known* parameter values, but the final result is presented as a substitute for the *uninformed* maximal expected payoff.

Several studies used stochastic sensitivity analyses to explore the parameter space more thoroughly by randomly drawing parameter values in relevant ranges. In [19], the authors computed average performances for different policies, but instead of averaging over all drawn parameter sets, they used a moving average along the values of one of the parameters. In an early article [5], the problem of parameter uncertainty could be swept aside by the authors as the same policy had the best performance for all drawn parameter values. This, however, needs not always be the case. In a more recent study [23], the authors presented their results in terms of the proportion of randomly drawn parameter settings for which each considered policy performed better than the others. Here we may raise the same objection as previously, that ranking policies in this way can be misleading as it somewhat mixes up informed and uninformed decision making. The authors also showed which policy was optimal in each region of the parameter space. Doing so undoubtedly gives insight into the problem hand and it may have significant practical implications for some specifications we will not tackle here. use a transparent metric in order to make the distinction between informed and uniformed decision makers and, for each, specify their best treatment strategies.

To support our claim that a clear distinction must be made between maximizing the expected performance and considering the expected optimal performance in analyses of empirical therapy policies, we will use numerical simulations of a variant of the compartmental models presented in [22], and specifically [7]. We will consider a family of adaptive empirical therapy policies in the same spirit as the policies presented in [16], and standard empirical therapy policies usually found in the literature. We will randomly draw parameter values and we will compute the performance of each policy given the drawn parameters. From there, we will be able to determine the policy with the best expected performance, and compare it with the expected performance of choosing the best policy for any parameter values. It is worth noticing that our estimates will depend on the ranges in which parameters will be drawn, and more specifically on our assumption that parameters are (for the most part) independent. The method we use in order to tackle uncertainty is Monte-Carlo sampling, hence we are confident that many generalizations – like parameters or individual-specific parameters – over our model would be handled without much technical issues.

Finally, we remark that the population dynamics parameters of the health care facility and of the disease are not the only sources of uncertainty relevant to the design of empirical therapy policies. Gathering information about the state of the epidemic, for instance, can be essential. The frequency of patient screenings has sometimes been interpreted as a quantity of information (e.g. in [21]). See also [15] for a dynamic optimization approach relying on periodic screening of the population, and a comparison with the expected performance of an optimization strategy [14] that does not use this information. Another source of uncertainty that is more difficult to control is the way policies are actually implemented in the field; see [4] for a stochastic differential equation approach to this problem. In the present article, we assume that the spread of the infection can be screened exactly when needed, and that the chosen policies are properly implemented.

We present the materials and methods used in the study in Section 2: the compartmental model describing the emergence and spread of an infection in a health care facility in Section 2.1, and the considered policies in Section 2.2. We show and discuss our results in Section 3: first in the uninformed case (Section 3.1), then in the informed case (Section 3.2). Section 4 concludes.

## 2 Materials and methods

### 2.1 Model

We use a compartmental epidemiological model to describe the spread of an infection in a health care facility (Figure 1). The events featured in the model are summarized in Table 1. The model parameters are shown in Table 2 with the distributions in which they are randomly drawn. We refer to [14] for a detailed description of the model, and for references supporting or bringing context to the underlying assumptions.

**Figure 1:**
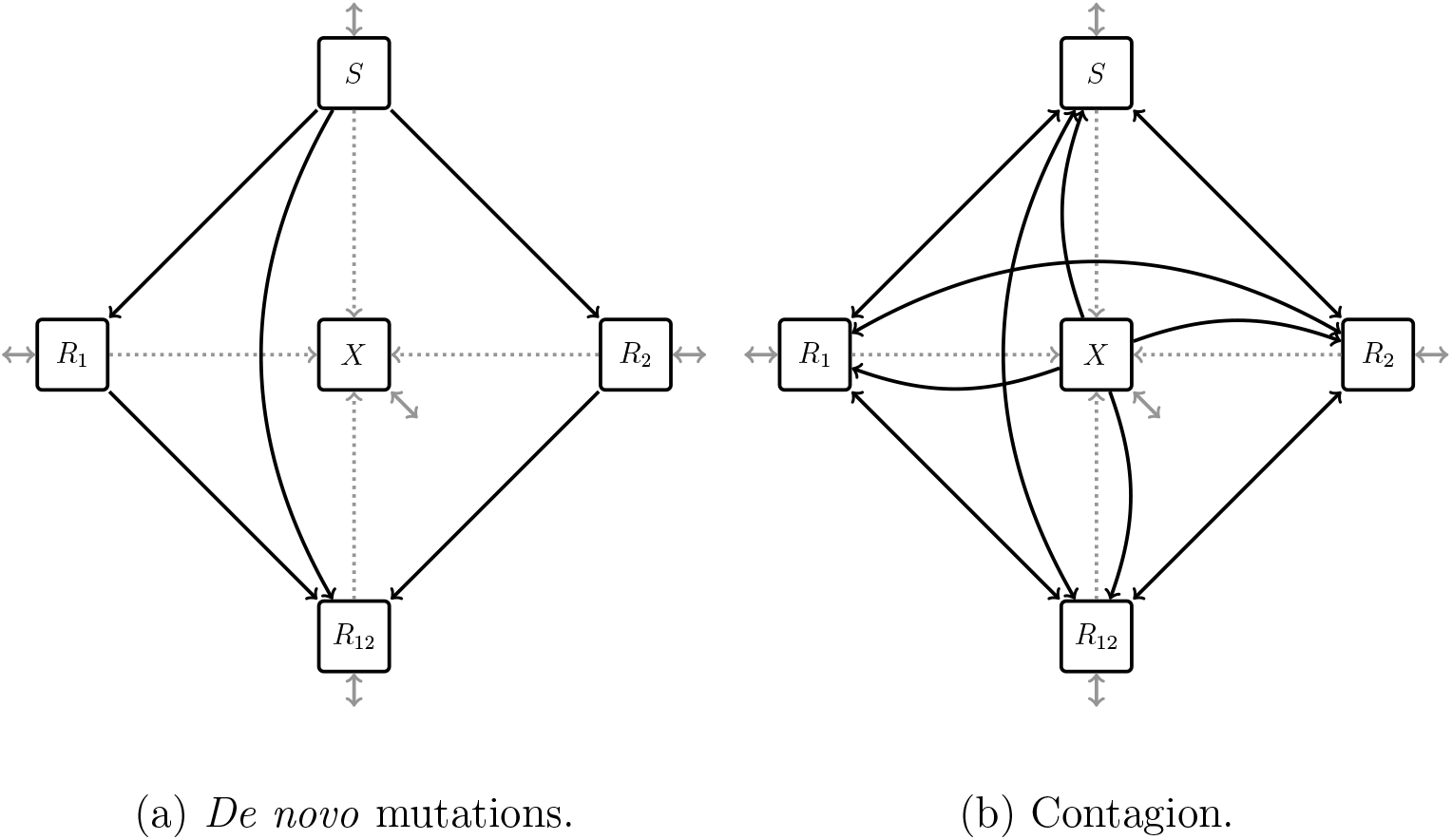
For clarity we display *de novo* mutations *(black, panel (a))* and contagion *(black, panel (b))* on two separate graphs. *Gray:* admissions and discharges. *Dotted gray:* recovery.

**Table 1:**
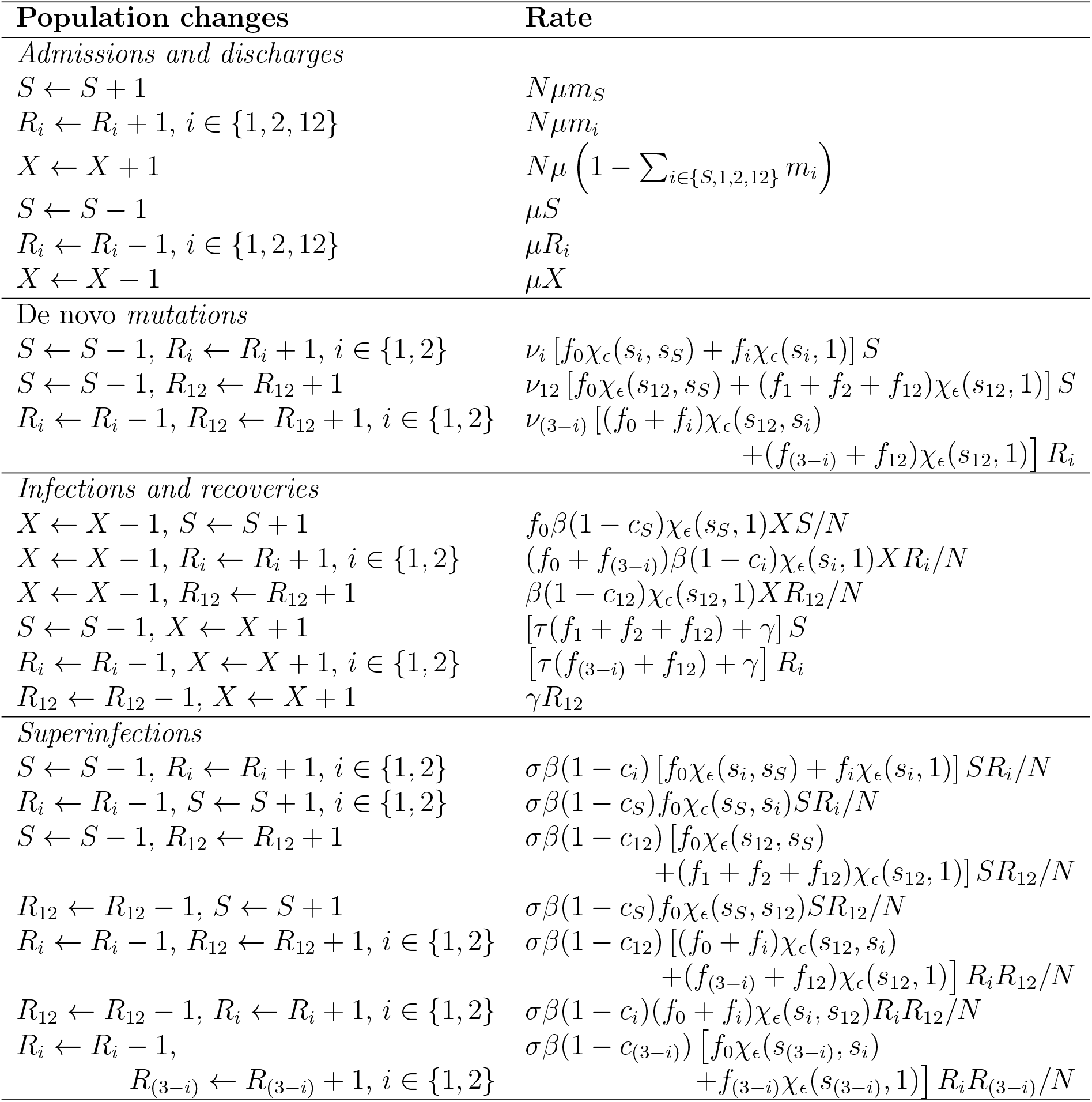
Events considered in the model.

**Table 2:** Model parameters. * source: [23].

| Parameter | Unit | Description | Range | Sampling |
| --- | --- | --- | --- | --- |
| *Admissions and discharges* |  |  |  |  |
| $N$ | – | Average number of patients. | $[10, 100]$ | linear |
| $\mu$ | day<sup>-1</sup> | Turnover rate. | $[3 \times 10^{-3}, 0.5]^*$ | logarithmic |
| $m_S$ | – | Proportion of incoming patients infected with the susceptible strain. | $[0.07, 0.9]^*$ | logarithmic |
| $m_1$ | – | Proportion of incoming patients infected with the strain resistant to drug 1. | $[10^{-3}, 0.1]^*$ | logarithmic |
| $m_2$ | – | Proportion of incoming patients infected with the strain resistant to drug 2. | $[10^{-3}, 0.1]^*$ | logarithmic |
| $m_{12}$ | – | Proportion of incoming patients infected with the strain resistant to drugs 1 and 2. | $[10^{-6}, 0.1]^*$ | logarithmic |
| *De novo mutations and fitness costs* |  |  |  |  |
| $\nu_i$ | day<sup>-1</sup> | Rate of acquisition of resistance to treatment $i$ through *de novo* mutation. $i \in \{1, 2, 12\}$ | $[10^{-10}, 0.1]^*$ | logarithmic |
| $c_i$ | – | Cost on infectivity of resistance to treatment $i$, $i \in \{1, 2, 12\}$. | $[0, 0.2]^*$ | linear |
| $c_S$ | – | Cost on infectivity of susceptible organisms. | 0 | – |
| $s_i$ | – | Cost on competitive performance of resistance to treatment $i$, $i \in \{1, 2, 12\}$. | $[0, 0.2]$ | linear |
| $s_S$ | – | Cost on competitive performance of susceptible organisms. | 0 | – |
| $s_X$ | – | Cost on competitive performance of commensal organisms. | 1 | – |
| $\epsilon$ | – | Slope parameter in function $\chi_\epsilon$. | $[0.05, 50]$ | logarithmic |
| *Infections, superinfections, and recoveries* |  |  |  |  |
| $\beta$ | day<sup>-1</sup> | Transmission rate. | $[0.2, 1]^*$ | linear |
| $\sigma$ | – | Relative rate of superinfection. | $[0, 0.5]^*$ | linear |
| $\gamma$ | day<sup>-1</sup> | Rate of spontaneous recovery. | $[0, 0.2]^*$ | linear |
| $\tau$ | day<sup>-1</sup> | Rate of recovery under appropriate treatment. | $[0.2, 1]$ | linear |

- *Pathogenic strains*. We model resistance to two drugs, drug 1 and drug 2. At each time, a patient is either infected with a wild-type strain susceptible to both drugs, infected with a strain resistant to drug 1 but not to drug 2 (1-resistant), infected with a strain resistant to drug 2 but not to drug 1 (2-resistant), infected with a strain resistant to both drugs (12-resistant), or uninfected with any of the above. *S*, *R*_1_, *R*_2_, *R*_12_, and *X* denote the number of patients with each health status respectively. We assume that infected patients are only infected with one pathogenic strain at any given time.
- *Admissions and discharges*. We consider a small health care facility or a hospital ward with an average population of *N* = 40 patients. We will run stochastic simulations of the model. We assume that admission and discharge do not depend on health status. We also make the simplifying assumption that transmission of strains and emergence of resistance in the community is exogenous to the problem in hand.
- *Treatments*. Three treatments are available: drug 1 in monotherapy (treatment 1), drug 2 in monotherapy (treatment 2), and a combination of drugs 1 and 2 (treatment 12). Patients can also be left without a treatment (treatment 0). Decision variable *f*_*i*_, *i* ∈ {0, 1, 2, 12}, is equal to 1 when patients receive treatment *i* and to 0 otherwise. At all time, *f*_0_ + *f*_1_ + *f*_2_ + *f*_12_ = 1.
- *Recovery*. Infected patients recover spontaneously at rate *γ* regardless of the infecting strain. Patients receiving adequate treatment recover at rate *τ*.
- De novo *mutation*. Infecting organisms acquire *i*-resistance through *de novo* mutation at rate *ν*_*i*_, *i* ∈ {1, 2, 12}, of two different genes as illustrated in Figure A.1 in Appendix.
- *Infection and superinfection*. We assume homogeneous mixing. Pathogenic strains are transmitted to uninfected patients with transmission rate *β*, and to already infected patients with transmission rate *σβ*. We assume *σ* = 1 in this instance of the model, but the general case is *σ ∈* [0,1]. *i*-resistant strains incur a fitness cost on transmission *c*_*i*_ ∈ [0,1], *i* ∈ {1, 2, 12}, and their transmission rate is weighed by 1 − *c*_*i*_.
- *Strain replacement*. Within-host, a new pathogenic strain acquired through *de novo* mutation or contagion must compete with the resident pathogenic strain or with the commensal microflora in the case of previously uninfected patients. *i*-resistant strains incur a fitness cost *s*_*i*_ ∈ [0,1], *i* ∈ {1,2, 12}, on competitive performance. We normalize the fitness cost *s*_*S*_ of the wild-type susceptible strain to 0, and the fitness cost *s*_*X*_ of the commensal microflora to 1. A strain receiving adequate treatment has a fitness cost of 1. The rate of replacement of a resident strain with fitness cost *s*_*j*_ by a new strain with fitness cost *s*_*i*_ is given by function *χ*_*ϵ*_ as

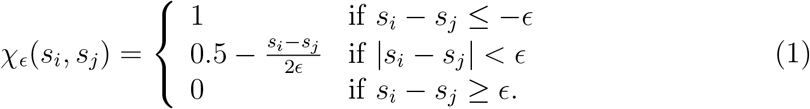

We further assume that a new strain cannot colonize a treated patient unless it is resistant to that patient’s treatment.

### 2.2 Empirical therapy policies

Our objective is to minimize the cumulative infected patient-days over two years (720 days). To do this, we assume that an empirical therapy policy is to be chosen among the THRES-*k*, *k* ∈ {0, …, 25}, adaptive policies defined as

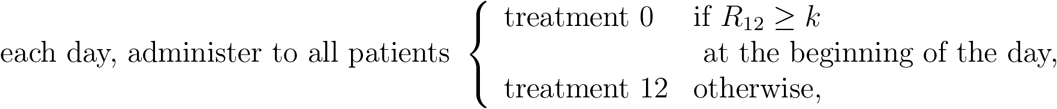

and THRES-*∞*. THRES-0 consists in administering treatment 0 to all patients during two years. THRES-*∞* consists in administering treatment 12 to all patients during two years; it corresponds to THRES-*k* in the limit *k* → ∞. Notice that in our setting, the threshold *k* does not depend on the average population size *N*.

There are several reasons why we focus on THRES-*k* empirical therapy policies in this study. First these policies have been shown to be close to policies optimized with artificial intelligence tools for some parameter values and when the number of individuals in each compartment is screened, see [15]. As we will see later in this article, they indeed yield much better results than standard policies. Also, THRES-*k* policies are much easier to implement in an actual health care facility. Finally, for our purpose, THRES-*k* policies are much less computationally demanding than actually optimized policies.

Combination therapies such as THRES-*∞* are frequently implemented in the clinic and are commonly considered in the literature. In the following discussions, we will at times compare the expected performances of the THRES-*k* polices to the expected performances of other policies found in the literature: MONO-1, MONO-2, CYC-*n* and METRO-*n*. MONO-1 and MONO-2 consist in administering treatment 1 and 2 respectively to all patients during two years. CYC-*n* is a cycling policy with a fixed period: treatment 1 during *n* days followed by treatment 2 during *n* days. METRO-*n* has been shown to be of special interest [14] and consists in administering treatment 12 during *n* days followed by no treatment during *n* days.

For the sake of clarification, let us make a some technical remarks at this point. In the present article, we are mainly concerned by the information the decision maker has or has not about the disease or population dynamics. When we say that the decision maker is informed or uninformed, we specify the status of the information about the disease and population dynamics parameters. However, the fact that we consider policies that are conditional – like THRES-*k* (*k* ∈ {1, …, 25}) policies – or unconditional – like THRES-*k* (*k* ∈ {0, ∞}), MONO-1, MONO-2, CYC-*n* and METRO-*n* policies – on the status of the individuals and in particular dependent on the number of individuals that are in compartment *R*_12_ is a different problem relying on a different type of information. Both sets of information are not exactly independent as we might learn the parameters of the disease or the population dynamics from the observations of the numbers of individuals in the different compartments at different times. We will not consider this possibility here and leave this aspect for further work.

## 3 Results

### 3.1 Uninformed decision making

Assuming that the exact values of the model parameters are unknown, we compare the policies presented in section 2.2 by randomly drawing *n*_*P*_ = 400 parameter sets. For each policy and each parameter set, we run *n*_*S*_ = 100 stochastic simulations of the model (section 2.1) and compute the average performance of the policy over *n*_*P*_ × *n*_*S*_ = 40, 000 stochastic simulations.^3^ All stochastic simulations are initialized by simulating a period of 30 years without treatment (treatment 0).

Under the THRES-0 policy, no treatment is ever implemented and we obtain an average of 27,920.4 cumulative infected patient-days over two years (95% CI: 26,345.6 – 29,495.2). The combination therapy THRES-*∞* allows an average 58.2% reduction compared to THRES-0 with an average cumulative infected patient-days: 11,668.9 (95% CI: 10,313.1 – 13,024.7). MONO-1 and MONO-2 yield respectively 20,356.2 (95% CI: 18,820.0 – 21,892.4) and 20,809.1 (95% CI: 19,249.1 – 22,369.1) cumulative infected patient-days on average, which corresponds to a 27.1% and a 25.5% reduction compared to THRES-0. The results obtained the tested CYC-*n* and METRO-*n* are not as good as the results obtained with THRES-*∞* (results shown in Table A.1 in the Appendix). Hence, of all the policies considered in this article that are also commonly found in the literature and that are independent on the number of individuals in the different compartments, THRES-*∞* shows the best performance; we retain THRES-*∞* as a benchmark in the following discussions.

The average performances of our adaptive policies THRES-*k* as a function of the threshold *k* are shown in Figure 2. We obtain the best performance with THRES-1, which consists in administering treatment 12 as long as no patient is infected with the 12-resistant strain, and treatment 0 when one or more patients are infected with the 12-resistant strain. Under THRES-1, the average cumulative infected patient-days is reduced by 74.7% compared to THRES-0 and by 39.3% compared to THRES-*∞* with an average of 7,078.1 (95% CI: 6,059.8 – 8,096.4) cumulative infected patient-days.

**Figure 2:**
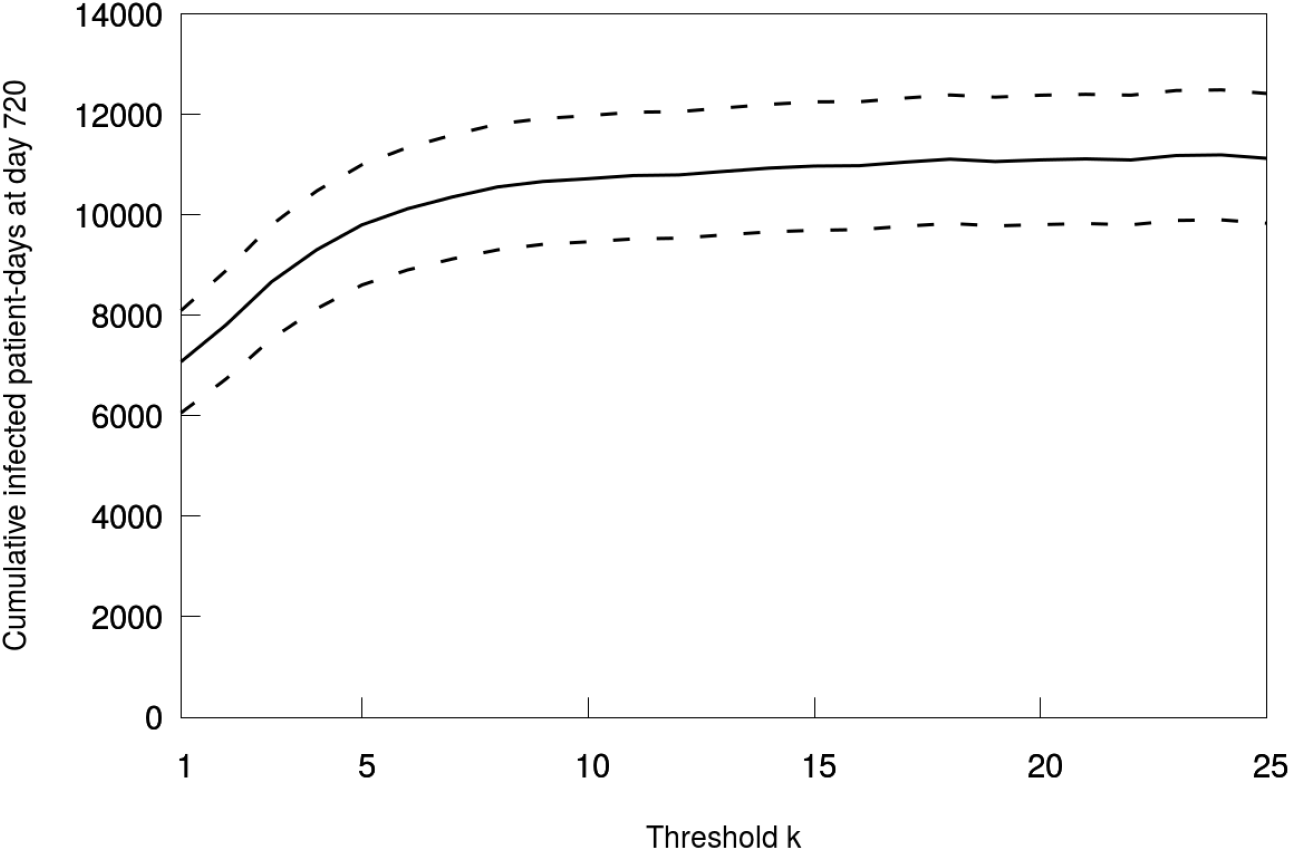
Average cumulative infected patient-days over two years (400 × 100 stochastic simulations) obtained with THRES-*k* policies. *Dashed:* 95% confidence intervals.

It is therefore reasonable for an uninformed decision maker to implement policy THRES-1 in order to minimize the expected cumulative infected patient-days over two years. THRES-1, however, will not show the best performance in all parameter settings. This is best illustrated in Figure 3: for each of the *n*_*P*_ = 400 randomly drawn parameter settings, we plot the average cumulative patient-days over *n*_*S*_ = 100 stochastic simulations obtained under THRES-*k* against that obtained under THRES-*∞*. We show zoomed-in graphs in Figure A.2 in Appendix. We see that THRES-*∞* outperforms THRES-1 for some parameter values (significantly at the 95% threshold). Unsurprisingly, the performance of THRES-25 is close to that of THRES-*∞*. Yet notice that we find a significant difference at the 95% threshold between THRES-25 and THRES-*∞* for several individual parameter settings.

**Figure 3:**
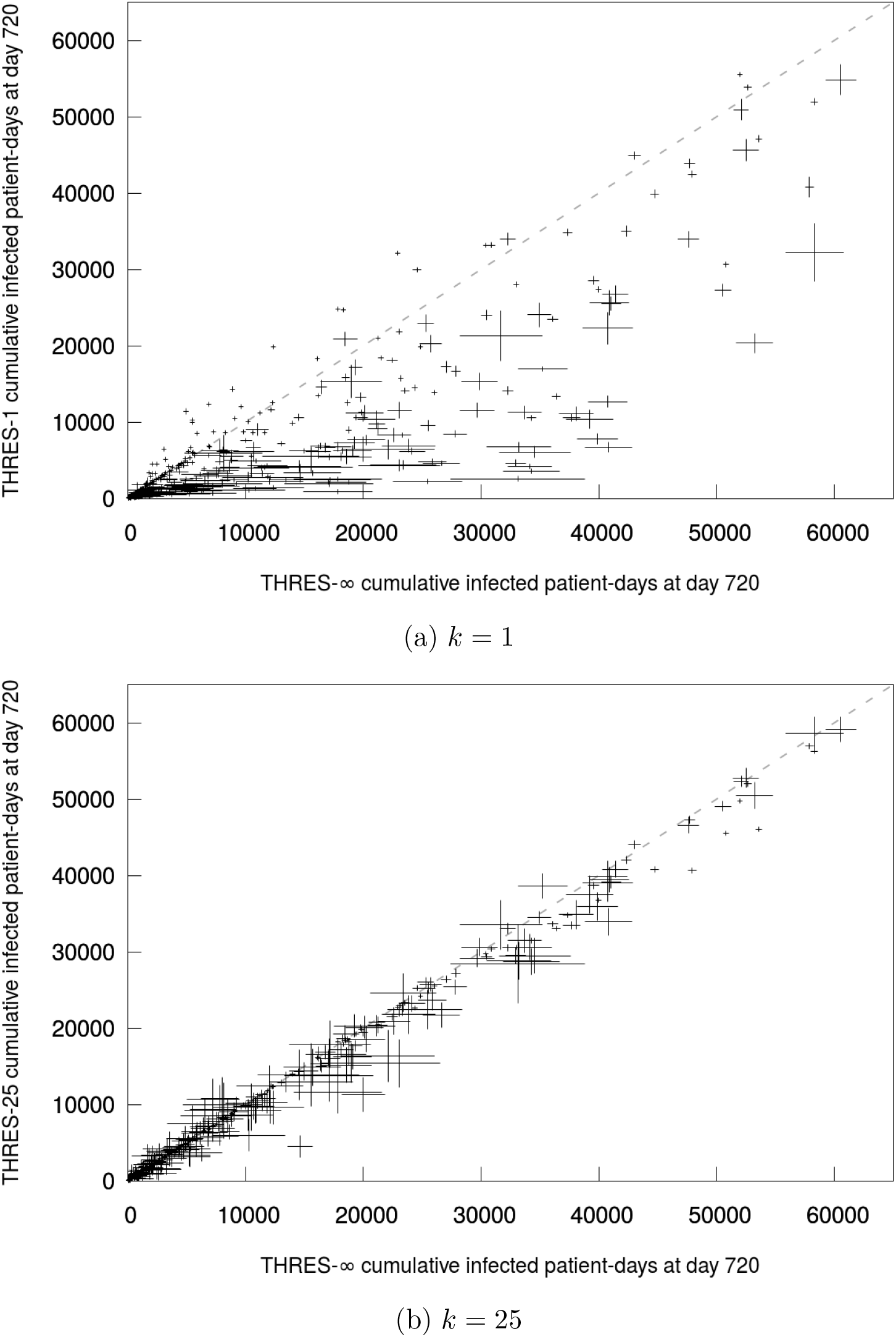
Average cumulative infected patient-days for THRES-*k* (*k* ∈ {1,25}) and THRES-∘. Each cross for one of 400 randomly drawn parameter settings. *Error bars:* 95% confidence intervals over 100 stochastic simulations. *Dashed:* y=x line.

### 3.2 Informed decision making

We now turn to decision making when the exact parameter values are known to the decision maker. In this setting, we call INFOBEST the empirical therapy policies chosen as follows. For each of the *n*_*P*_ = 400 randomly drawn parameter settings, we compute the average two year cumulative infected patient-days over *n*_*S*_ = 100 stochastic simulations for each considered policy (THRES-*k*, *k* ∈ {0..25}, and THRES-*∞*), and we retain the policy with the best average performance. Put differently, INFOBEST policies maximize the expected performance given known parameters.

Then we run another series of *n*_*S*_ stochastic simulations under the chosen INFOBEST policy for each parameter setting. Notice that we use different runs for the same strategies in order to avoid statistical biases not differentiating between the training runs used to define our policy and the test runs which outcomes correspond to our *in silico* trials. By definition, the average performance of the INFOBEST policies over the *n*_*P*_ ×*n*_*S*_ simulations is better than any of the THRES-*k* (*k* ∈ {0, …, 25, ∞}) and in particular THRES-1, the policy maximizing expected performance in an uninformed setting. We obtain an average difference of 278.3 (95% CI: 147.0 – 409.6) cumulative infected patient-days over two years between INFOBEST and THRES-1. INFOBEST allows a 3.9% improvement over THRES-1. This difference estimates the expected value of knowing the exact value of population dynamics and disease parameters. The average cumulative infected patient-days obtained with INFOBEST for each parameter value are plotted against those obtained with in Figure 4.^4^

**Figure 4:**
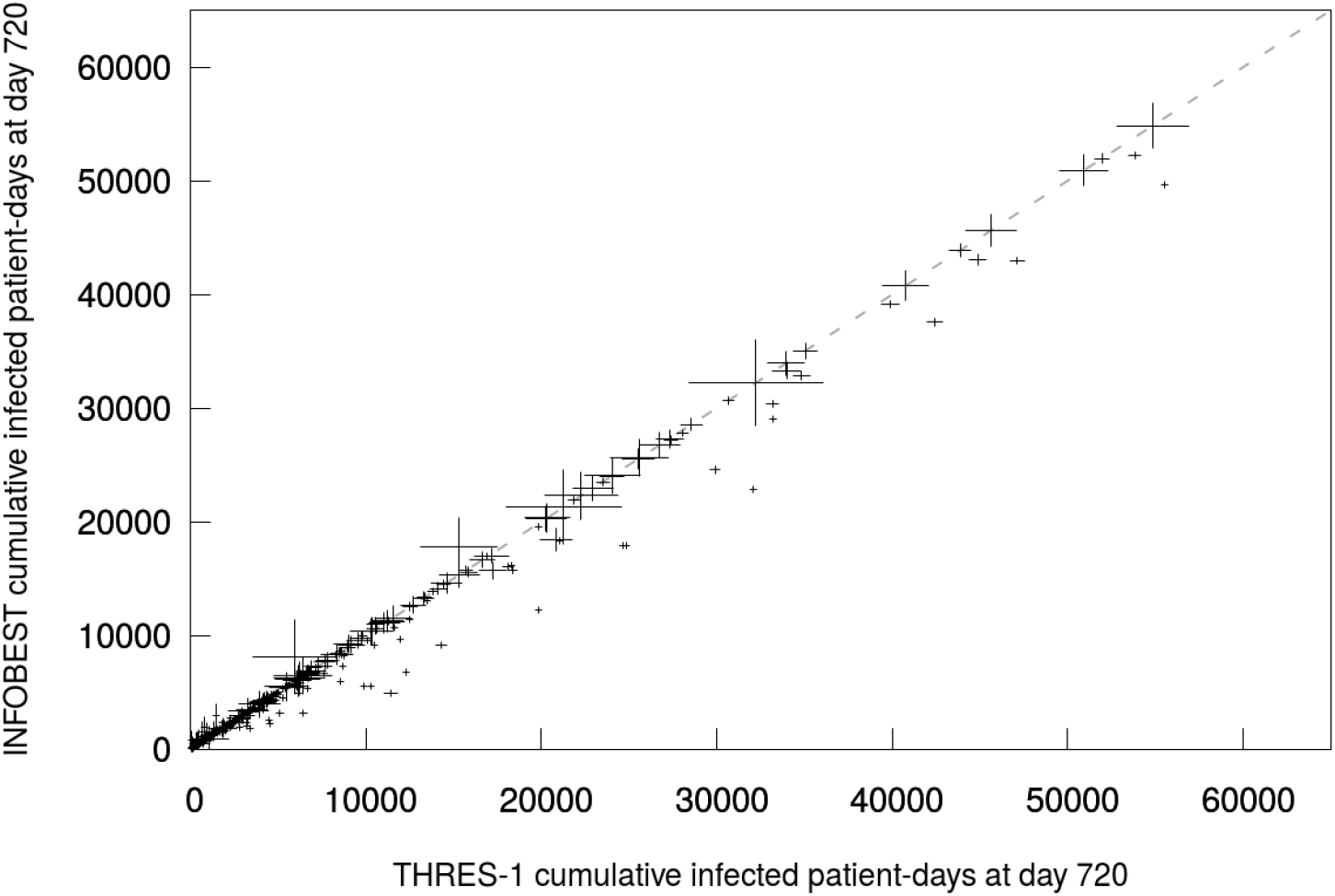
Average cumulative infected patient-days for INFOBEST and. Each cross for one of 400 randomly drawn parameter settings. *Error bars:* 95% confidence intervals over 100 stochastic simulations. *Dashed:* y=x line.

Our comparisons of policies have relied so far on the average cumulative infected patient-days. It must be noted that our analysis did not internalize the costs associated with the different policies, for instance screening costs and treatment costs. To be complete in this setting, costs must be considered once a policy has been chosen. In Figure 5, we show the average proportion of days with treatment 12 over two years obtained with the THRES-*k* and the INFOBEST policies. For thresholds *k* ≥ 8, the proportion of days in which treatment 12 is administered is increasing with *k*, as the threshold *k* above which all treatment is stopped becomes less likely to be crossed. Counter intuitively, the average proportion of days with treatment 12 is decreasing with *k* for *k* ≤ 7; THRES-1, the policy that would be chosen in the uninformed case, is among the policies implying the most days with treatment 12. This shows that small thresholds are efficient at reducing the emergence of double resistance, thus allowing to administer treatment 12 more often overall. The average proportion of days with treatment 12 obtained with the informed INFOBEST policies is higher (0.833, 95% CI: 0.830 – 0.836) than those obtained with the THRES-*k* policies for *k* ∈ {1, …, 25}. Indeed, for some parameter settings, the INFOBEST policy is THRES-*∞* for which treatment 12 is administered everyday. In particular, the average proportion of days with treatment 12 is significantly lower at the 95% threshold under THRES-1 (0.791, 95% CI: 0.788 – 0.794) than under INFOBEST. This may have policy implications, as it shows that if treatment costs are not internalized in the analysis, informed decision making increases the expected drug consumption compared to uninformed decision making.

**Figure 5:**
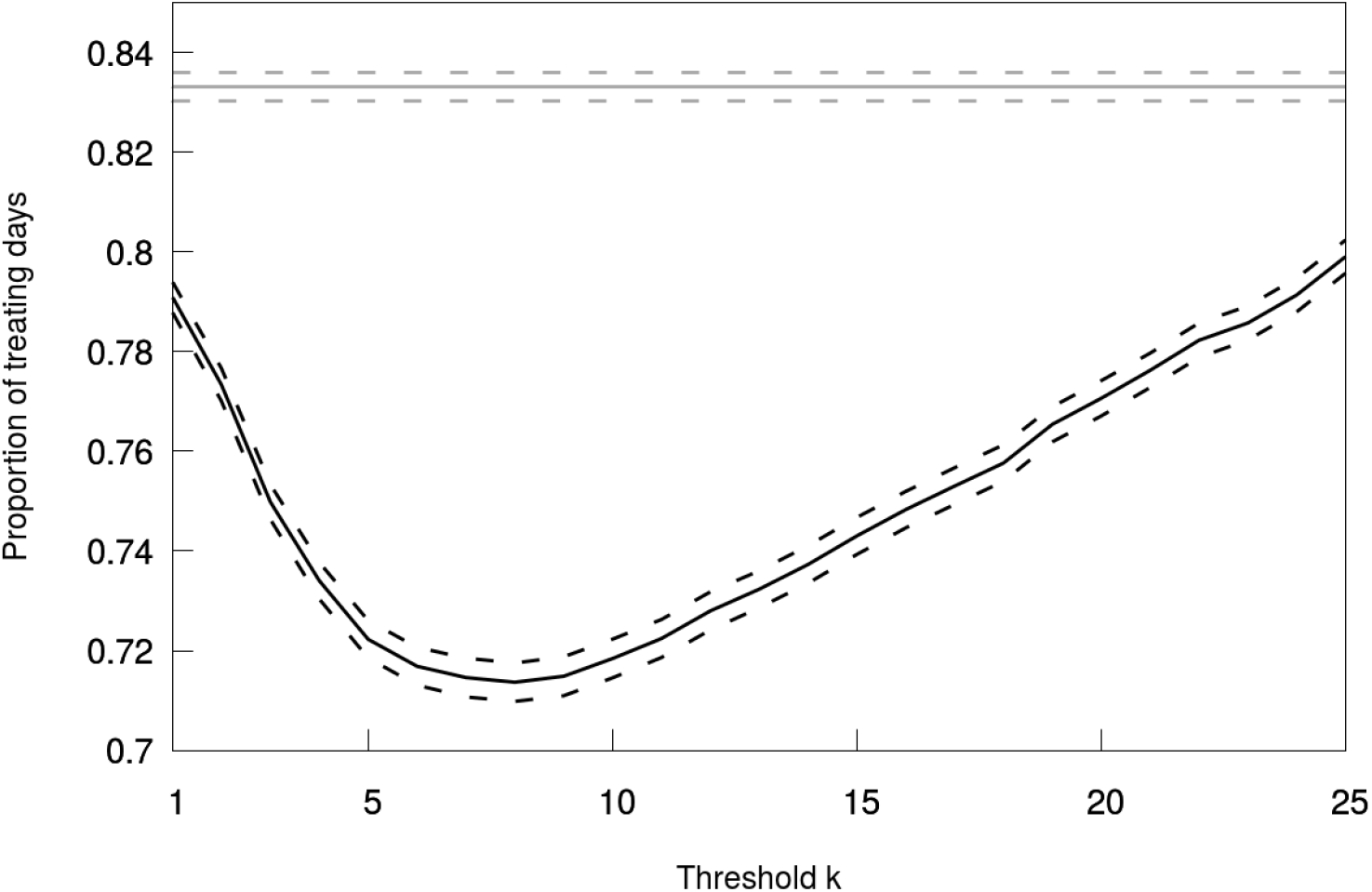
Average proportion of days with treatment 12 over two years (400 × 100 stochastic simulations). *Black:* THRES-*k*. *Gray:* INFOBEST (independent on *k*). *Dashed:* 95% confidence intervals.

## 4 Conclusion

An essential element in the design of empirical therapy policies is the information available to the decision maker. Being able, for instance, to set screening priorities, is an important part of empirical therapy devising and analysis. Yet doing so requires to make a proper distinction between informed and uninformed decision making when producing and presenting results.

In this article, we focus on the information regarding the population dynamics parameters of the disease and of the considered health care facility. We emphasize the distinction between decision making in uninformed and informed settings. objective is to minimize the cumulative infected patient-days in the long run (two years). In an uninformed setting, only the range of each parameter and the underlying dynamical system are known to the decision maker. An empirical therapy policy is then chosen that minimizes the expected cumulative infected patient-days over two years. In an informed setting, the exact parameter values are known, and the decision maker chooses a policy that minimizes the expected cumulative infected patient days (over stochastic realizations of the spread of the disease) given the parameter values.

To illustrate our point, we draw upon the stochastic version of a compartmental epidemiological model describing the spread of an infecting organisms in a health care facility. The organism may evolve resistance to two antimicrobial drugs. We assume that a policy is to be chosen in a family of adaptive empirical therapies that consist in administering a combination therapy when the number of patients infected with a double resistant strain is below a threshold, and to administer no drug when this threshold is exceeded. The combination therapy is a limiting case of such adaptive policies.

In the uninformed setting, we find that the best adaptive policy reduces the average cumulative infected patient-days over two years by 39.3% (95% CI: 30.3% – 48.1%) compared to the combination therapy. Choosing an empirical therapy policy knowing the exact parameter values allows to avoid a further improvement of 3.9% (95% CI: 2.1% – 5.8%). This estimates the expected value of having access to exact parameter values. While this figure may seem only marginal, we point out that it is still informative: it shows that gathering more information about parameter values may not be cost efficient, at least in the setting presented in this article. Indeed, we also show that drug consumption is higher on average in the informed setting. A full consideration of the costs associated with the policies might therefore shift the balance towards uninformed expected performance maximization.

## A Additional figures and results

**Figure A.1:**
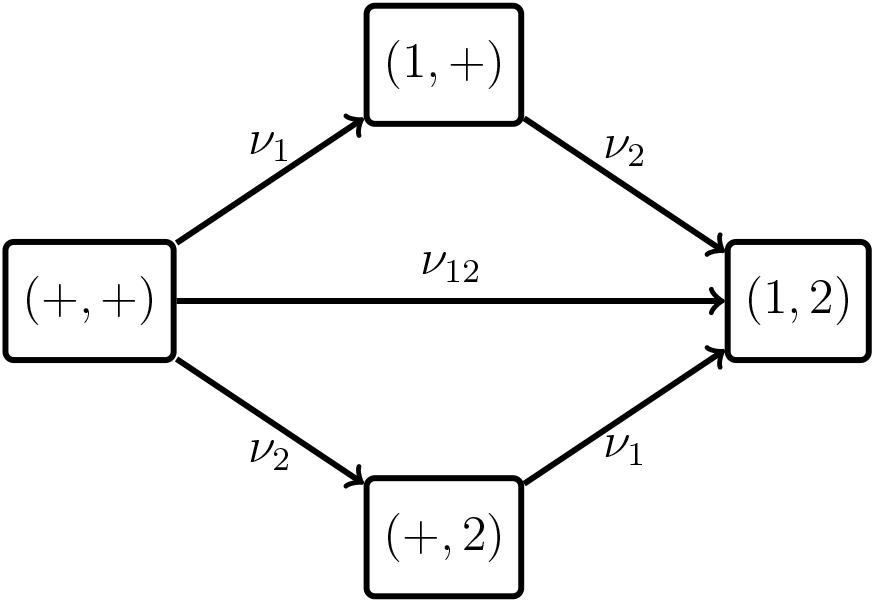
The four genotypes included in our model with mutation rates. The arrows represent transitions between genotypes. The two considered genes are separated by a comma. ‘+’ represents a wild-type allele. ‘1’ and ‘2’ represent alleles giving resistance to drugs 1 and 2 respectively. (+, +): wild-type susceptible. (1, +): 1-resistance only. (+, 2): 2-resistance only. (1, 2): resistance to both drugs.

**Table A.1:** Outcomes for CYC-*n* and METRO-*n* and different values of *n*.

| Policy | Cumulative infected patient-days at day 720 | Policy | Cumulative infected patient-days at day 720 |
| --- | --- | --- | --- |
| CYC-10 | 13,353.4<br>(95% CI: 11,979.9 – 14,726.9) | METRO-10 | 15,536.8<br>(95% CI: 14,210.5 – 16,863.1) |
| CYC-20 | 14,265.5<br>(95% CI: 12,886.0 – 15,645.1) | METRO-20 | 16,172.1<br>(95% CI: 14,912.4 – 17,431.8) |
| CYC-30 | 14,855.6<br>(95% CI: 13,481.0 – 16,230.1) | METRO-30 | 16,471.6<br>(95% CI: 15,249.8 – 17,693.5) |
| CYC-40 | 15,399.4<br>(95% CI: 14,018.2 – 16,780.7) | METRO-40 | 16,656.5<br>(95% CI: 15,449.9 – 17,863.1) |
| CYC-50 | 15,707.3<br>(95% CI: 14,328.4 – 17,086.2) | METRO-50 | 16,601.0<br>(95% CI: 15,414.4 – 17,787.6) |
| CYC-60 | 16,046.7<br>(95% CI: 14,658.2 – 17,435.2) | METRO-60 | 16,938.5<br>(95% CI: 15,752.3 – 18,124.6) |

**Figure A.2:**
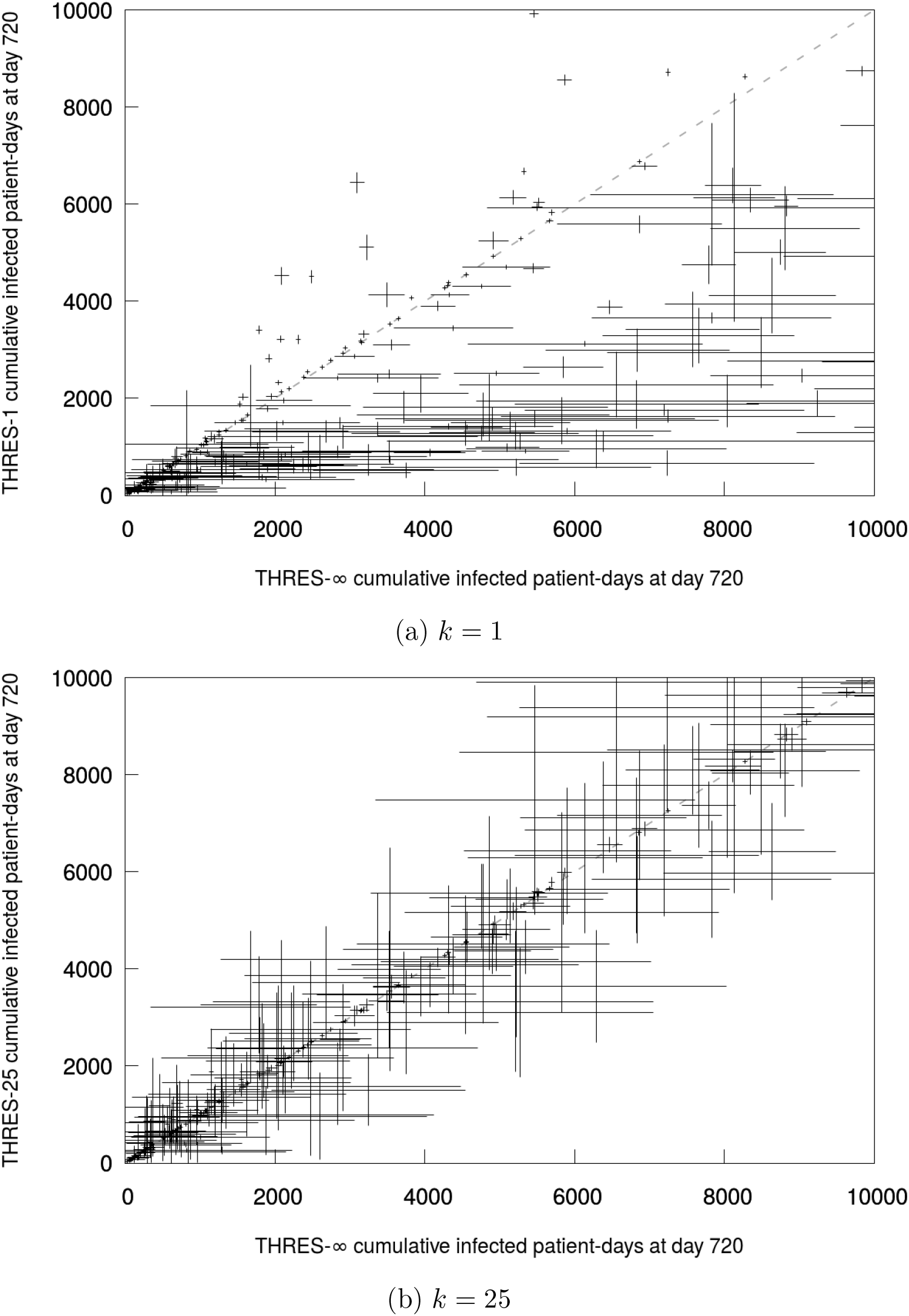
Average cumulative infected patient-days for THRES-*k* (*k* ∈ {1, 25}) and THRES-∞. Each cross for one of 400 randomly drawn parameter settings. *Error bars:* 95% confidence intervals over 100 stochastic simulations. *Dashed:* y=x line. Same as Figure 3 with different scales.

**Figure A.3:**
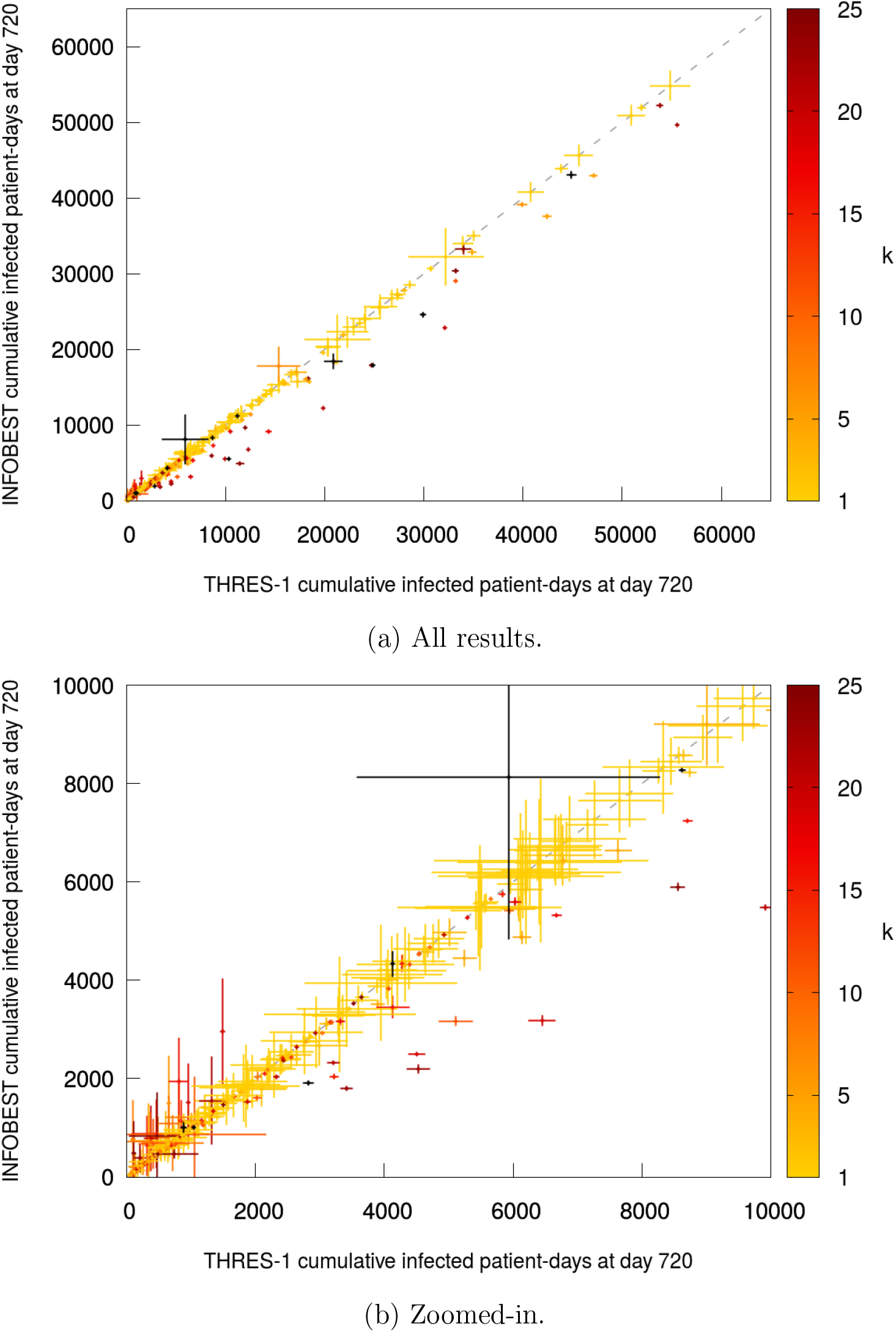
Average cumulative infected patient-days for INFOBEST and. Each cross for one of 400 randomly drawn parameter settings. *Error bars:* 95% confidence intervals over 100 stochastic simulations. *Dashed:* y=x line. Color corresponds to the implemented strategy that maximizes outcome: yellow-red for THRES-*k* for *k* ∈ {1..25}, black for THRES-*∞*.

## Footnotes

1 In the following, we will include prophylactic treatments in this definition.

2 For the sake of simplicity, in this presentation we use the expected value but any other aggregation function could be applied similarly.

3 The confidence intervals of the grand means (means of means) are computed with the appropriate random effects.

4 Alternative views are provided in Appendix.

